# Comprehensive machine learning boosts structure-based virtual screening for PARP1 inhibitors

**DOI:** 10.1101/2024.03.15.585277

**Authors:** Klaudia Caba, Viet-Khoa Tran-Nguyen, Taufiq Rahman, Pedro J. Ballester

## Abstract

Poly ADP-ribose polymerase 1 (PARP1) is an attractive therapeutic target for cancer treatment. Machine-learning scoring functions constitute a promising approach to discovering novel PARP1 inhibitors. Cutting-edge PARP1-specific machine-learning scoring functions were investigated using semi-synthetic training data from docking activity-labelled molecules: known PARP1 inhibitors, hard-to-discriminate decoys property-matched to them with generative graph neural networks and confirmed inactives. We further made test sets harder by including only molecules dissimilar to those in the training set. Comprehensive analysis of these datasets using five supervised learning algorithms, and protein-ligand fingerprints extracted from docking poses and ligand only features revealed two highly predictive scoring functions. The PARP1-specific support vector machine-based regressor, when employing PLEC fingerprints, achieved a high Normalized Enrichment Factor at the top 1% on the hardest test set (NEF1% = 0.588, median of 10 repetitions), and was more predictive than any other investigated scoring function, especially the classical scoring function employed as baseline.

**Scientific Contribution:** We present the first PARP1-specific machine-learning scoring functions for structure-based virtual screening. A particularly rigorous evaluation, including test sets with novel molecules and a much higher proportion of challenging property-matched decoys, reveals the most predictive scoring function for this important therapeutic target. Typically, narrow machine learning analyses would have likely missed this promising PARP1-specific scoring function, which is now released with this paper so that others can use it for prospective virtual screening.

**Key Points:**

- A new scoring tool based on machine-learning was developed to predict PARP1 inhibitors for potential cancer treatment.
- The majority of PARP1-specific machine-learning models performed better than generic and classical scoring functions.
- Augmenting the training set with ligand-only Morgan fingerprint features generally resulted in better performing models, but not for the best models where no further improvement was observed.
- Employing protein-ligand-extracted fingerprints as molecular descriptors led to the best-performing and most-efficient model for predicting PARP1 inhibitors.
- Deep learning performed poorly on this target in comparison with the simpler ML models.

## Introduction

PARP1 plays an important role in regulating the microhomology-mediated end joining (MMEJ) pathway that repairs DNA damage [1,2]. PARP1 is a member of the diphtheria toxin-like ADP-ribosyltransferases family that is catalytically activated in response to various types of DNA damage [1,2]. The full-length PARP1 protein is modular and composed of six domains (Figure 1) [3]. In the N-terminus, the first three domains are zinc finger domains (Zn1, Zn2 and Zn3), followed by a BRCT (BRCA1 C-terminus domain) and a WGR (Trp-Gly-Arg) domain. At the C-terminus, there is a catalytic domain (CAT), which houses a helical domain (HD) and an ADP-ribosyl transferase (ART) domain. These domains allosterically communicate with each other in order to facilitate DNA damage repair. The three zinc finger domains at the N-terminus recognize both DNA single- and double-strand breaks, thereby causing conformation changes in the CAT domain that allow NAD^+^ to be recognized at the activation site en route to activating the enzyme. Activated PARP1 catalyzes the poly ADP-ribosylation of susceptible protein substrates using NAD^+^ [4,5]. Furthermore, PARP1 has also been found to play a role in transcriptional regulation, chromosome stability, cell division, differentiation, apoptosis, and has been considered the most actively-pursued target for treating some cancers [6]. Breast Cancer Type 1 Susceptibility Protein (BRCA1) and Breast Cancer Type 2 Susceptibility Protein (BRCA2) have a crucial function in DNA damage repair via homologous recombination pathway. In cells with deleterious BRCA mutations, the MMEJ pathway becomes critical for the repair of DNA damage. As such, inhibition of PARP1 by NAD^+^ competitive inhibitors can prevent the repair of DNA damage in BRCA-deficient cancer cells, leading to cancer cell apoptosis [5,7]. PARP1 is thus a validated therapeutic target for ovarian and/or breast cancer with deleterious BRCA mutations [8].

**Figure 1.**
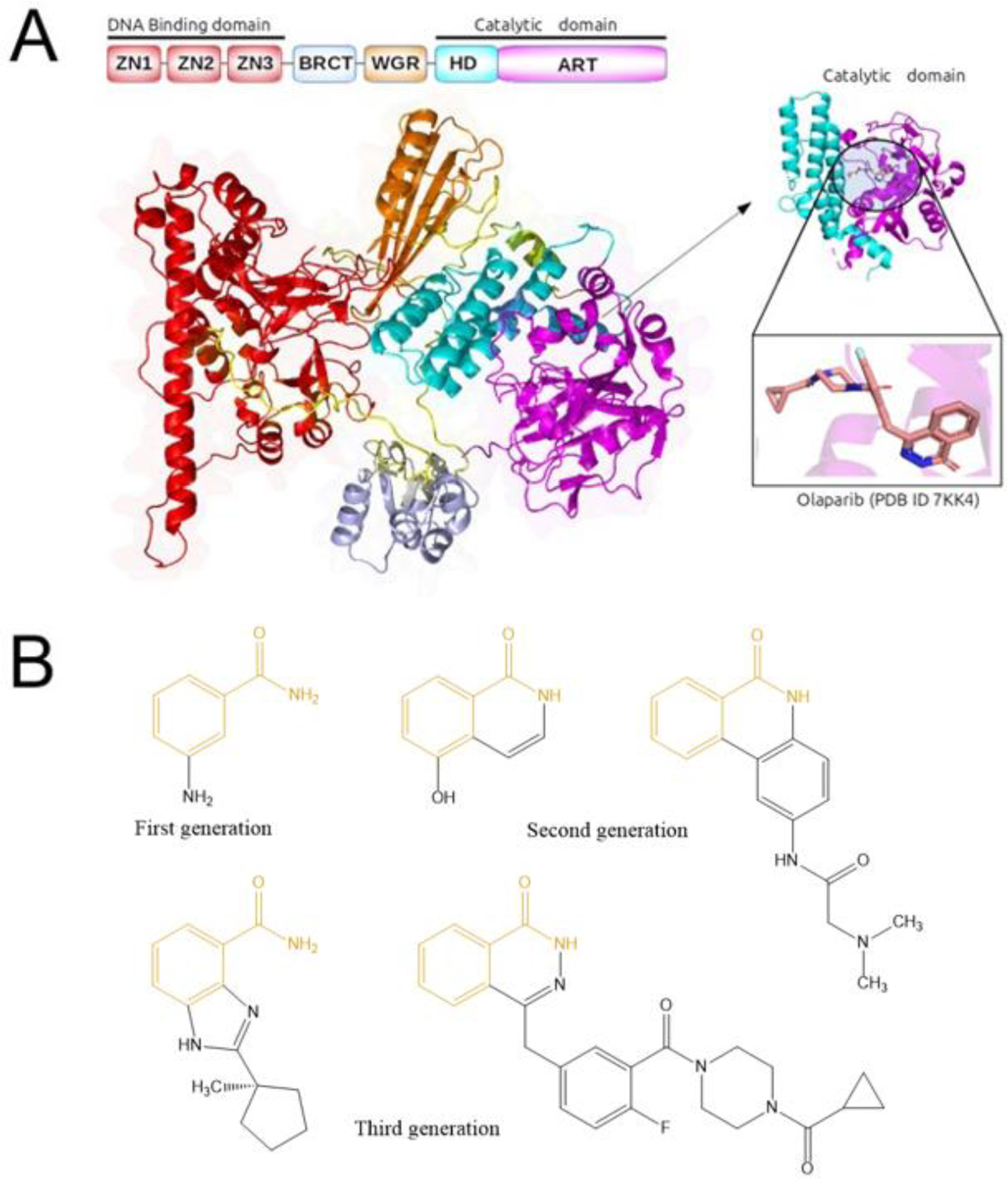
PARP1 and its inhibitors. (A) Cartoon illustration of the PARP1 protein. Catalytic domain co-crystalized with ligand (Olaparib) was based on the protein structure of PARP1 (PDB code 7KK4). (B) Structures of several PARP1 inhibitors from three generations (Olaparib is the third-generation inhibitor on the right). Benzamide scaffold common to all FDA-approved PARP1 inhibitors shown in yellow.

Small-molecule PARP1 inhibitors have become the standard of care for women with metastatic ovarian cancer and breast cancer harbouring the single or double BRCA1 and BRCA2 mutations [9,10]. These inhibitors can be categorized into three major types according to their discovery timelines [11]. First-generation PARP1 inhibitors are benzamide derivatives (e.g. 3-aminobenzamide) and close analogues of nicotinamide-related compounds [12]. They were discovered via empirical drug design, given that benzamide was the first to show an inhibitory activity against PARP1 [12], and nicotinamide, a by-product of the PARP1 enzymatic reaction, is a weak PARP1 inhibitor [11]. Second-generation inhibitors contain a quinoline core and were first reported in 1991 [11,13,14]. These PARP1 inhibitors were shown to slow down the repair of DNA damage [14], and were later optimized to enhance their potency using structure-based drug design. This helped improve the understanding of PARP1’s active site and facilitated the synthesis of highly potent novel inhibitors (around 50 times more potent than 3-aminobenzamide) [15]. Third-generation inhibitors were the first to show that PARP1 inhibitors can exert their activity when used alone in BRCA-mutated cancer patients. Widely regarded as a major breakthrough in PARP1-related cancer research [16,17], this discovery represented a paradigm shift in cancer treatment that triggered an intense period of activity centered on the development of PARP1 inhibitors, leading to the approval of olaparib, rucaparib, niraparib, and talazoparib by the FDA [11]. All FDA-approved PARP1 inhibitors contain the benzamide scaffold [18]. These drugs do not work in combination with first-line chemotherapy, due to additive hematological toxicity [19]. Frequent delivery of olaparib, niraparib and rucaparib may result in toxic accumulation in normal tissues due to poor absorption and distribution [20]. Other drawbacks have also been recorded, e.g., non-selective PARP1 activity, low solubility and low permeability, as in the case of olaparib [21]. Therefore, the discovery of novel PARP1 inhibitors is still highly sought after [19,22].

Structure-based (SB) virtual screening (VS) has been shown to be useful in hit identification for a range of therapeutic targets [23]. The available atomic-resolution structures of PARP1 and the affinities/activities of their cognate ligands support the use of docking to predict both protein-ligand binding affinity and the plausible binding modes of an inhibitor to the CAT of PARP1. SBVS relies primarily on molecular docking, which entails two main challenges. In the first step, the correct pose of the molecule must be sampled (sampling), and the second step involves ranking and selection of the correct pose (scoring). Many methods exist for rapidly exploring the conformational space of a small molecule [24,25]. The latter step is used to guide the sampling process as well as to rank the sampled poses. Putative pose scores can be used to direct the sampling strategy, as implemented in AutoDock Vina [26], or to analyze the fitness of an entire population, as in AutoDock4 [27]. In any scenario, an accurate scoring function is essential for ranking and selecting samples.

As non-linear relationships between the chosen protein-ligand features and the binding affinity/bioactivity of the ligand should not be neglected, there is a need for methods that will take such complex relationships into account. Machine learning (ML)-powered docking does take into account nonlinearities [28,29] and has discovered molecules with affinity for a range of targets [30–36]. Docked molecules are ranked according to their predicted affinity/energy of interactions for the target using a scoring function (SF). The more accurate SF predictions are, the more tightly-bound compounds should be placed at the top of the post-screening hit list. Although useful in some cases, hit ranking by classical SFs, which rely on linear assumptions is generally quite limited [37]. A target-specific ML SF is a computational model employed to rescore docked poses that effectively leverages the data available for the investigated target [38].

Here, we will investigate SB models to predict PARP1 inhibitory activity via target-specific ML SFs aiming at identifying the most potent molecules. We will report a retrospective SBVS study with training data derived from different experimental settings. We will compare the models based on data sets relevant to PARP1 and three off-the-shelf generic SFs including Smina, CNN-Score, and SCORCH. In this paper, we will also discuss the impact of different modelling choices on VS performance on PARP1 as well as the benefit of using regression-based ML SFs trained on inactive-enriched data for SBVS.

## Results

### Experimental design

The experimental design of this study is illustrated in Figure 2. A data set of compounds tested against PARP1 and their inhibitory activity (potency/affinity) values were obtained from ChEMBL (Version 29), consisting of 5097 bioactivity data points, which belong to different categories including half maximal inhibitory concentrations (IC_50_s) from biochemical or cellular assays (biochemical/cellular IC_50_s), or binding affinity (K_d_ values) from biophysical assays and others. Compounds capable of inhibiting PARP1 at a concentration lower than or equal to 1 µM were classified as actives, while those having an active concentration above 1 µM were labeled as inactives. The threshold of 1 µM is a usual choice in the literature [39–42], consistent with what can be found in bioactivity databases (e.g., in PubChem, nearly 90% of PARP1 inhibitors have a potency value better than 1 µM, whereas all inactive-labeled compounds cannot inhibit this target at a concentration lower than this threshold [43]). Chemical structures of PARP1 inhibitors were processed according to a previously described data curation workflow [44], which starts by the removal of metal ions and salts, the normalization of chemotypes and the standardization of tautomers using the JChem Standardizer. Four PARP1 structures in the Protein Data Bank (PDB) having good resolutions (1.50-2.10 Å) were considered (PDB IDs: 7AAC, 7KK5, 7KK4, 6VKK), and 7KK4 was selected for this study. Olaparib (CHEMBL521686), a clinical PARP1 inhibitor, was co-crystallized in 7KK4 structure. The data set was divided into two subsets: the training set and the test set.

**Figure 2.**
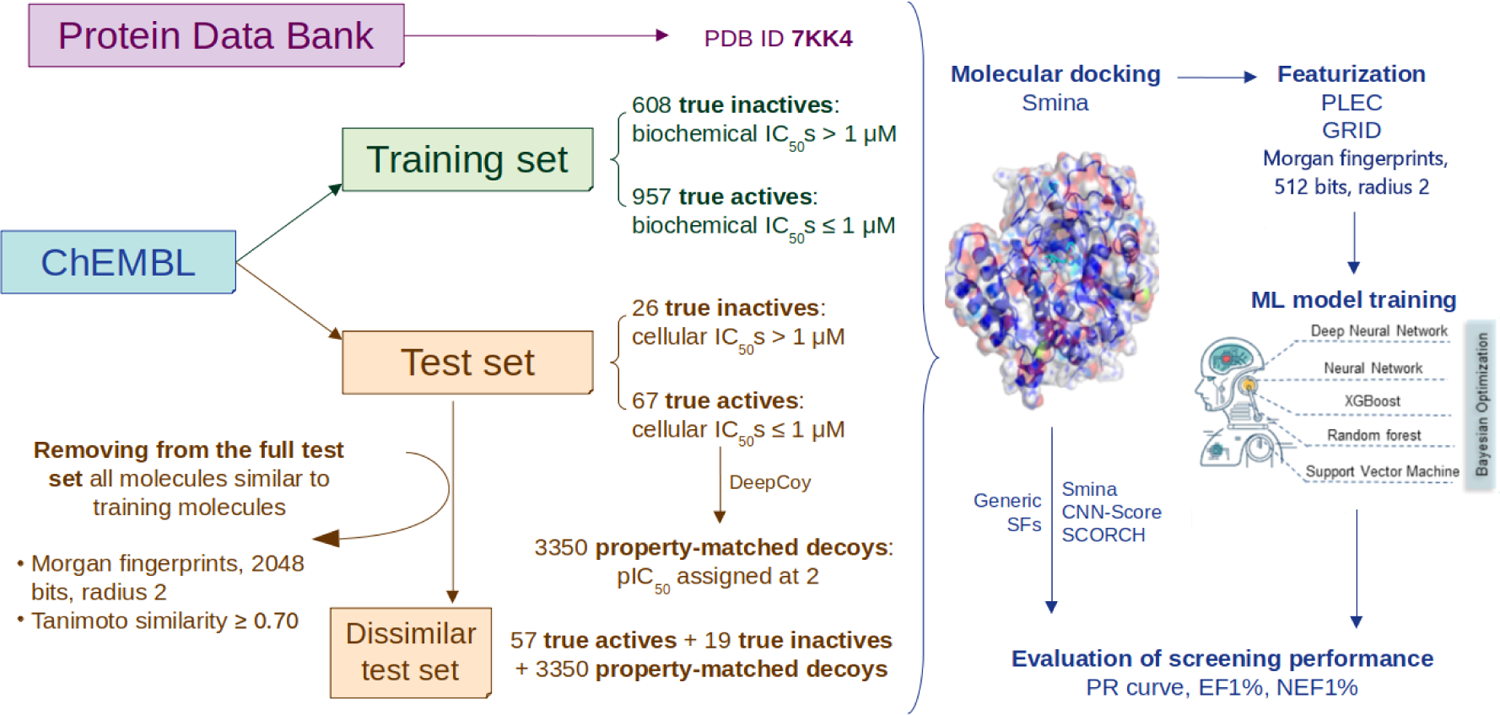
The methodological workflow of this study. Experimental data on PARP1 were retrieved from ChEMBL and distributed to the training set and the test set. The training set contains 1565 molecules with biochemical (but not cellular) IC_50_s, while the test set comprises 93 molecules with cellular IC_50_s in addition to their *in vitro* biochemical potency. The threshold to distinguish true actives from true inactives in both data sets is 1 µM. 3350 decoys property-matched to 67 test actives were generated by DeepCoy, and form part of the test set. All ligands were docked into their receptor (PDB ID 7KK4) by Smina, after which either PLEC or GRID features were extracted from each docked complex. Ligand-only Morgan fingerprints (512 bits, radius 2) were also computed. These features were then used as input for ML model training and testing, using five supervised learning algorithms: RF, SVM, XGB, ANN, DNN (hyperparameters were tuned using Bayesian optimization). The VS performance of all algorithms was evaluated in terms of EF1% and NEF1% and visualized as precision-recall curves. Three off-the-shelf generic SFs (Smina, CNN-Score, SCORCH) were evaluated on the same test set as the ML SFs for comparison. A dissimilar test set was also created by keeping only test molecules whose Tanimoto similarity scores to any training instances (Morgan fingerprints, 2048 bits, radius 2) were lower than 0.70, on which all SFs were also evaluated.

Activity records were specifically filtered to keep only values supplied with a standard relation type as “=” (certain data) or “>” (classified as inactives). Molecules with the sought cellular activity are important in that many molecules with on-target activity are discarded because of not being cell active and hence data is partitioned accordingly. The training set consists of 957 molecules with biochemical (but not cellular) IC_50_s (certain data) not exceeding 1 µM, classified as actives, and 608 compounds whose active concentrations were above 1 µM, classified as inactives. By contrast, the test set comprises 93 molecules with cellular IC_50_s (certain data) in addition to their *in vitro* biochemical potencies, 67 of which are actives (cellular IC_50_s ≤ 1 µM). DeepCoy next used these test actives as templates to generate property-matched decoys (decoy-to-active ratio = 1:50), outputting 3350 decoys which form part of the test set. This is a graph-generative neural network that creates new decoys in an iterative and bond-by-bond manner, such that they are chemically similar but structurally dissimilar to their input active. An additional difficulty posed by this PARP1 benchmark is that test inactives (decoys) are related to their test actives in a different way from training inactives to training actives. This avoids performance overestimation caused by decoy bias, which occurs when training and test inactives are both property-matched to training and test actives, respectively [45]. Moreover, the test set was made even more challenging for SBVS, by removing all test molecules whose Tanimoto similarity scores to any training instances (Morgan fingerprints, 2048 bits, radius 2) were equal to 0.70 or above. This dissimilar test set aims at examining the discriminatory power of each ML SF on molecules structurally dissimilar to its training data. We first introduced this more demanding evaluation of SFs in a recent study [39].

Five supervised learning algorithms, each with a binary classification variant and a regression variant, were used to develop our ML SFs: random forest (RF) [46], extreme gradient boosting (XGB) [47], support vector machine (SVM) [48], artificial neural network (ANN) [49], and deep neural network (DNN) [50]. These algorithms have been employed in many studies to train high-performing ML models for *in silico* screening in drug discovery, and were featured in a recently-published protocol [39]. Here, they were trained using different featurization schemes, including protein-ligand complex-based features (protein-ligand extended connectivity fingerprints, PLEC; or 3D grid-based features, GRID) [39,51], and ligand-only Morgan fingerprints (512 bits, radius 2) [52], which are commonly used for encoding structural information carried in a target-ligand complex or a ligand structure in 3D space. The VS performance of all PARP1-specific ML SFs on the full and dissimilar test set versions was then compared to that achieved with off-the-shelf generic ones (Smina, SCORCH and CNN-Score) [53–55]. We plotted the precision-recall (PR) curve, which shows the trade-off between the precision and recall values at different cutoffs of the ranked list of test molecules, for each SF. We also computed the enrichment factor in the top 1%-ranked test molecules (EF1%) and the normalized EF1% (NEF1%), two useful metrics for evaluating how well each SF retrieves true actives among its top-ranked compounds (early enrichment), which is an important aspect in VS, notably when the SF is used in prospective settings [39,56]. The EF1% is computed as the hit rate in the top 1%-ranked molecules divided by that across the whole library of compounds, indicating how many times more actives are retrieved among the top 1%-ranked molecules by a certain SF than by random guessing [39]. The NEF1%, on the other hand, is the EF1% recorded for an SF divided by the maximal EF1% it can possibly achieve on a given test set. This metric permits comparing the VS performance of multiple SFs on the same test set, and also across test sets (notably those of different sizes) [39,57].

### Applying existing generic SFs

Smina, a fork of Autodock Vina (version 1.1.2), was used to dock all molecules in this study (https://sourceforge.net/projects/smina/files/). The native SF of Smina is a linear regression model trained on the CSAR-NRC HiQ 2010 data set. This SF is used to predict the binding free energy of a docked pose [53]. The utilization of customized interaction terms, together with high-quality training data sets, improves its predictive performance over the original AutoDock Vina SF. The native SFs from ten other docking programs including Dock4, DockIt, FlexX, Flo+, Fred, Glide, Gold, LigandFit, MOEDock and MVP were less successful than Smina at distinguishing active compounds from pharmacologically relevant decoys. These data contain a large number of closely related compounds for which experimental affinities have been measured using a standard protocol for a diverse set of targets including serine/threonine protein kinase, trypsin-like serine protease, bacterial type II topoisomerase, methionyl tRNA synthetase, hepatitis C RNA polymerase, polypeptide deformylase from *E. coli*, polypeptide deformylase from *S. Pneumococcus*, and peroxisome proliferator-activated receptor [25].

Several studies reported that rescoring the docked poses of the screened molecules resulted in better VS performance than relying solely on classical SFs used by docking programs [58]. For this purpose, two other off-the-shelf generic SFs were evaluated in this study. First, CNN-Score takes comprehensive 3D representations of a protein-ligand interaction as input and uses deep ML techniques to rescore docked poses [59]. It is an ensemble of five convolutional neural network (CNN) models of deep learning architecture (up to 20 hidden network layers). Of these models, the ‘dense’ and ‘default2017’ CNN models were trained using a large proportion of assumed inactives, in particular property-matched decoys extracted from the Database of Useful Decoys: Enhanced (DUD-E) [60], containing 22,645 positive instances and 1,407,145 negative instances. It should be noted that PARP1 is one of the 102 targets of DUD-E, and only 18 molecules used to train CNN-Score’s underlying models (out of 3443, i.e. 0.52%) were included in the test set: this represents a tiny overlap between the training set and the test set that would unlikely result in overestimating the screening power of this SF. CNN-Score was shown to perform better than two classical SFs (Smina and Vinardo) on LIT-PCBA [61]. Second, SCORCH consists of three ML approaches (gradient-boosted decision trees, feedforward neural networks and wide-and-deep neural networks). These ML SFs were trained on the PDBbind data set (refined set of 4854 complexes), the Binding MOAD data set (non-redundant set of 3187 complexes), the Iridium data set (highly trustworthy set of 120 complexes), and property-matched decoys generated from the DeepCoy generator with a decoy-to-active ratio of 10. SCORCH was proven better-performing than widely used SFs in both VS and pose prediction scenarios on independent data sets [54].

The VS performance of these three generic SFs is reported in Table S1, with a graphical illustration in Figure 3. Among them, the ML-based ones (CNN-Score and SCORCH) outperformed the classical SF (Smina), in terms of (N)EF1%. This suggests that deep learning and consensus models with largely available training data could help increase the screening power of SFs in SBVS campaigns. The advantage of generic SFs is that they can be used off-the-shelf to rescore docked poses issued by any docking tool in a relatively fast and simple manner [60].

**Figure 3.**
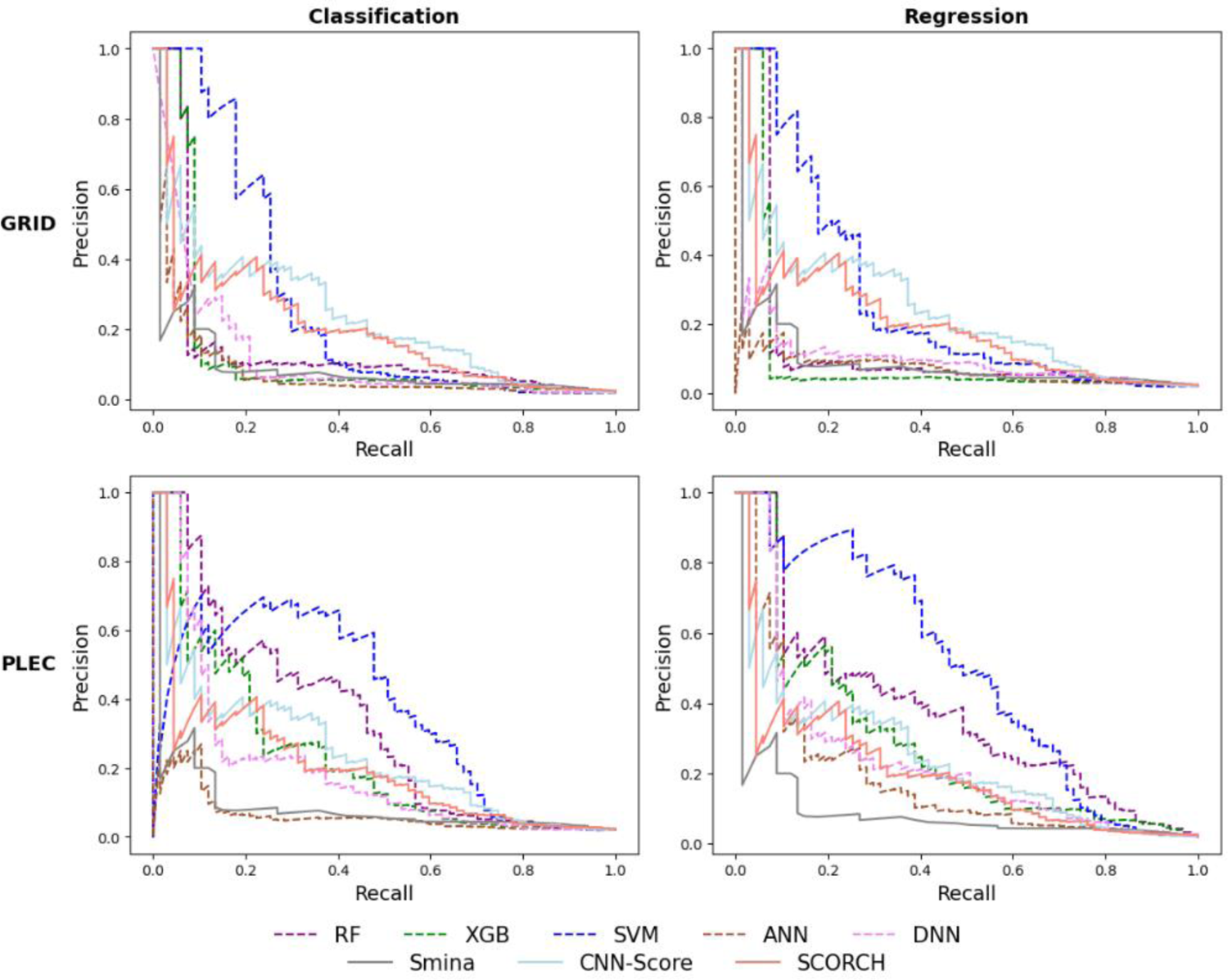
Precision-recall curves given by the generic and PARP1-specific SFs. To generate PARP1-specific ML SFs, docked poses of the PARP1-ligand complex were encoded either as GRID features (top) or as PLEC fingerprints (bottom). The resulting features were used by each of the following classification (left) and regression (right) learning algorithms: RF (purple, dashed line), XGB (green, dashed line), SVM (blue, dashed line), ANN (sienna, dashed line), and DNN (violet, dashed line). The PR curve of each target-specific ML SF is that of the training-test run giving an NEF1% equal (or closest) to the median NEF1% across 10 runs (chosen at random if multiple runs satisfy this criterion). The generic SFs are represented as solid lines in gray (Smina), light blue (CNN-Score), and salmon (SCORCH). Results are further specified in Table S1.

### Developing PARP1-specific models for SBVS using protein-ligand features

A way to improve VS performance further is to develop target-specific SFs [62]. The lack of confounding factors due to molecular recognition differences from other targets (e.g. those related to protein structures) means that a more accurate features-activity relationship can often be determined [63]. Target-specific SFs trained with a relatively low proportion of inactive molecules tended not to perform well on SBVS [64–66], and so it was avoided here. Furthermore, such SFs are designed to find active compounds in a chemolibrary containing a much higher percentage of inactives. This prompted us to evaluate the PARP1-specific ML SFs using a test set enriched with inactives. Their VS performance was summarized in Table S1 and depicted in Figure 3.

Target-specific classification and regression models built on PLEC fingerprints were found to be more predictive than those based on GRID features: the areas under the PR curves (PR-AUCs) of the SFs issued from the former type of features are generally larger than those produced by the latter type (Figure 3). The PLEC-based SF employing SVM as learning algorithm performed better than any other classifier and all three generic SFs on the test set. Indeed, this model retrieved over 32 times more true actives among the top 1%-ranked molecules than what would be expected at random. In the case of regression-based SFs, the majority of those trained with PLEC features gave better performance than their classification counterparts (except for RF regressor which performed worse than the RF classifier). In particular, the combination of PLEC and SVM, as featurization scheme and learning algorithm, once again led to the best performance: EF1% = 38.8, NEF1% = 0.764. This further demonstrates the importance of using quantitative bioactivity data points to train PARP1-specific ML SFs.

### Investigating the impact of combining features on training and testing ML SFs

We would like to investigate whether the use of ligand-only features alongside protein-ligand ones would result in better ML model training. For this purpose, Morgan fingerprint features, also known as extended-connectivity fingerprints ECFP4 [67], were employed. These features have frequently been used in binding affinity prediction among various topological parameters [68]. By combining them with protein-ligand features (PLEC, as it led to better-performing models than GRID), additional models were generated. The VS performance of these models is indicated in Table S2 and depicted in Figure 4. It is observed that the best-performing ML SF trained on the combination of Morgan fingerprints and PLEC features is again the regression-based SVM model, which achieves the largest PR-AUC on our test set (Figure 4). This SF performed equally well as the SVM regressor trained on PLEC features alone, in terms of early enrichment of test actives (NEF1% = 0.764). As the latter requires generating only one set of features, it is more efficient and there is no benefit in introducing ligand-only features in this case. The combination of Morgan fingerprints and PLEC features did, however, result in RF, XGB, SVM classifiers and RF, XGB, ANN and DNN regressors with more discriminatory power than their counterparts trained on PLEC features only, as evidenced by their PR-AUCs portrayed in Figure 4 and their (N)EF1% values indicated in Table S2.

**Figure 4.**
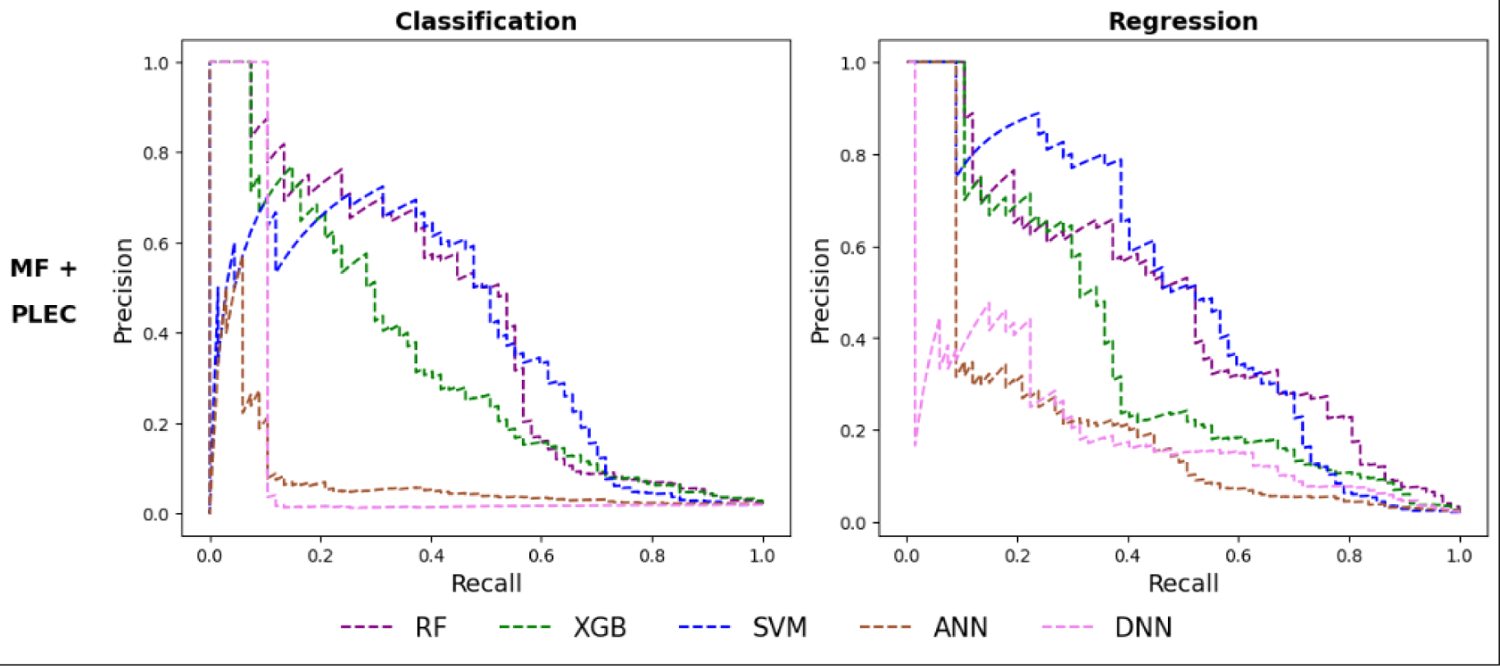
Precision-recall curves given by the PARP1-specific SFs based on combined Morgan fingerprints and PLEC features. To generate PARP1-specific ML SFs, docked poses of the PARP1-ligand complex were encoded as Morgan fingerprint (MF) features combined with PLEC fingerprints. The resulting features were used by each of the following classification (left) and regression (right) learning algorithms: RF (purple, dashed line), XGB (green, dashed line), SVM (blue, dashed line), ANN (sienna, dashed line), and DNN (violet, dashed line). The PR curve of each target-specific ML SF is that of the training-test run giving an NEF1% equal (or closest) to the median NEF1% across 10 runs (chosen at random if multiple runs satisfy this criterion). Further details are provided in Table S2.

### VS performance on the dissimilar test set

The employed test sets are hard in that each active has a high number of decoy molecules with highly similar physico-chemical properties. We now make it even harder by removing any test molecule similar to at least one training molecule. More concretely, a dissimilar test set was generated from the full test set, by discarding all test molecules having Tanimoto similarity scores ≥ 0.70 to any training molecule, in terms of Morgan fingerprints (2048 bits, radius 2), as introduced in a recently-developed protocol [39]. This way, the remaining test data are dissimilar to the training set, and are expected to be more challenging for VS, as structural biases in the training-test composition are reduced. Here we examined the target-specific ML SFs employing PLEC fingerprints, or a combination of Morgan fingerprints and PLEC fingerprints as features, because these two featurization schemes performed the best on the full test set (Tables S1, S2; Figures 3, 4). The NEF1% values of all SFs, both generic and target-specific (classifiers and regressors) were computed for the dissimilar test set and compared to those obtained from the full test set (Figure 5, Table S3).

**Figure 5.**
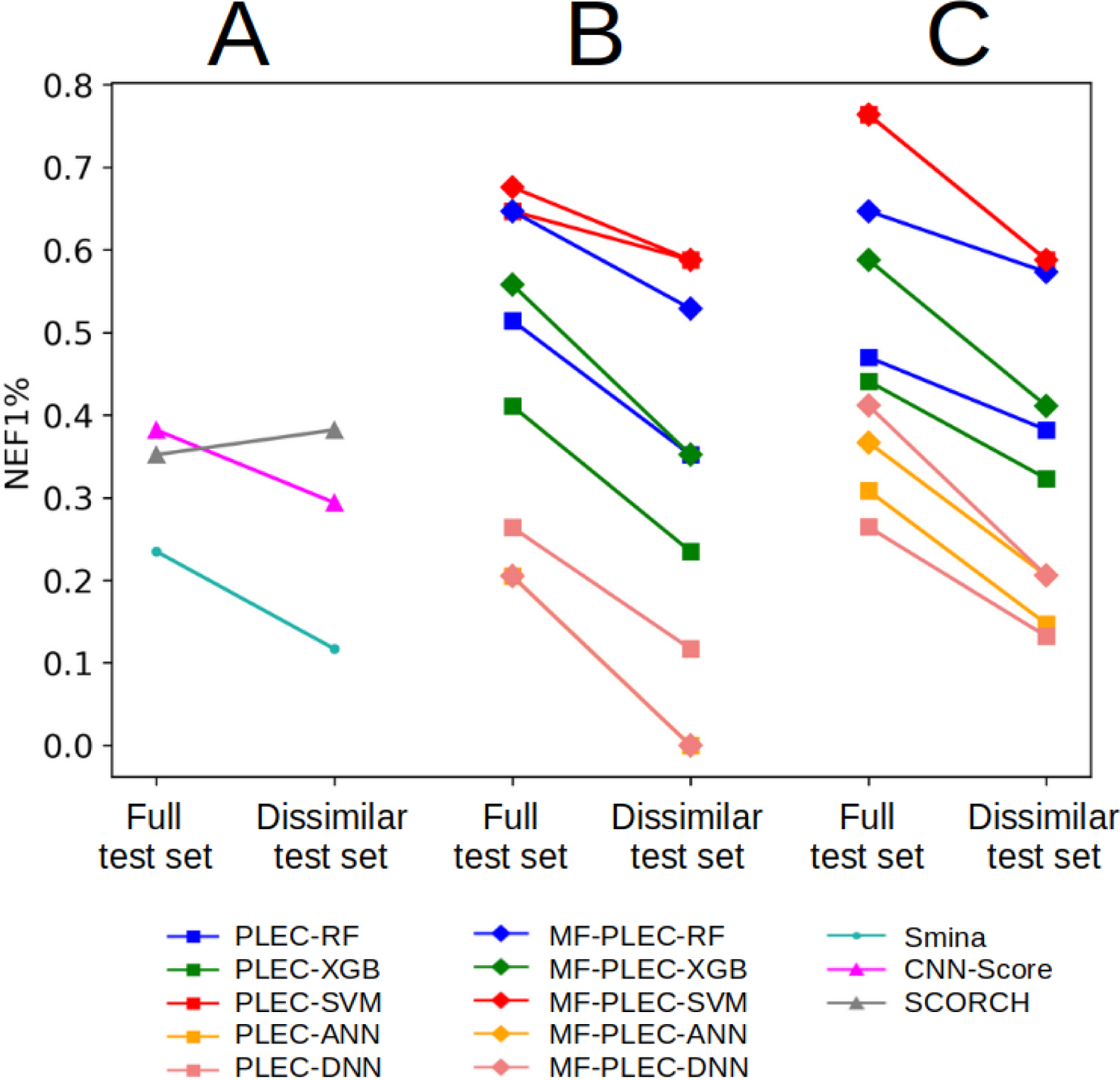
Screening performance of SFs on the full and dissimilar test set versions in terms of NEF1%. The SFs are: generic SFs **(A)**, classification-based target-specific ML SFs **(B)**, and regression-based target-specific ML SFs **(C)**. The NEF1% of each ML SF-test set pair is the median obtained after 10 training-test runs of the corresponding learning algorithm on the respective test set. Random NEF1% values for the full and dissimilar test set versions are 0.020 and 0.017, respectively.

It can be observed in Figure 5 that the dissimilar test set is indeed more challenging than the full test set: most SFs (22 out of 23, 95.65%) obtained a lower NEF1% on the dissimilar test set than on its full version (the only exception is SCORCH, whose performance was better on the dissimilar version of the test set: Tables S1, S3). 16 out of 20 target-specific ML SFs still performed better than Smina (except for classification-based ANN and DNN models trained on either PLEC only or a combination of Morgan fingerprints and PLEC features). Notably, the four ML SFs that performed the best on the full test set (the SVM-based regressor and the SVM-based classifier using either PLEC only or Morgan fingerprints and PLEC) still gave the best performance on the dissimilar test set (their NEF1% was 0.588, much higher than those of other SFs, including generic ones). These observations suggest that the exclusion of test molecules structurally similar to training data did not impact the relative comparisons of the investigated SFs in terms of VS performance. On a side note, SCORCH, a recently introduced generic ML SF, performed worse than eight PARP1-specific ML models on the dissimilar test set. Four of them outperformed SCORCH by retrieving 53.85% more true actives in the top 1%, showing the importance of building target-specific SFs whenever possible (results on PARP1 suggest that it is worth considering SCORCH when there are few known inhibitors for the target).

It is also worth noting that the similarity cutoff of 0.70 in terms of Morgan fingerprints already resulted in challenging test sets, as acknowledged in a previous study [39]. Here, the performance of the evaluated SFs dropped, in nearly all cases, when they were applied to our dissimilar test set (generated with this cutoff, Figure 5). Even though the maximal similarity is allowed to reach 0.70 for at least one pair of training-test molecules, the average similarity across all training-test pairs is only 0.18. Also, results with other ligand similarity cutoffs in a closely related problem showed that the performance for intermediate cutoffs, including 0.70, was pretty similar [37].

### The impact of computational modeling choices on SBVS performance

The boxplots in Figure 6 demonstrate the distributions of NEF1% values obtained from the test set according to: (i) the nature of the SFs: classification or regression (A); and (ii) the featurization scheme: GRID or PLEC (B), PLEC alone or in combination with Morgan fingerprints (C). The Shapiro-Wilk test was first carried out to assess the normality of the NEF1% data, giving a p-value < 0.05, implying that the NEF1% achieved by the SFs were not normally distributed. Welch’s t-tests were thus used to examine whether any two compared groups listed above were significantly different from each other, in terms of NEF1%.

Figure 5 shows that the performance of ML SFs on PARP1, despite varying widely depending on the modeling choices, was generally well above that of the best classical SF. This is typically the case across other targets [33,35,39,66,69], and hence we explained that comparing a classical SF with just a few ML SFs is misleading [69]. For example, the best-performing SVM regressor trained on PLEC features strongly surpassed Smina in terms of discriminatory power (NEF1% of 0.764 versus 0.235, respectively, Table S1). Moreover, out of the 20 ML SFs in Figures 5B and 5C, only the two ANN classifiers and a DNN classifier (15%) obtained a slightly worse NEF1% than Smina (0.205 versus 0.235 on the full test set, Tables S1 and S2; 0.000 versus 0.117 on the dissimilar test set, Table S3). In this context, should anyone only compare these three SFs to Smina, they would incorrectly conclude that ML SFs were generally not better than classical ones on this target.

Interestingly, as VS results from all featurization schemes were taken into account, the overall performance of regressors was not significantly better than that of binary classifiers on this target (Figure 6A). As seen in Figure 6B, GRID features clearly led to poorer-performing models on the full test set than those trained with PLEC fingerprints. The models employing the latter features, in turn, gave significantly poorer VS performance than those trained with both PLEC and Morgan fingerprint features in combination (Figure 6C).

**Figure 6.**
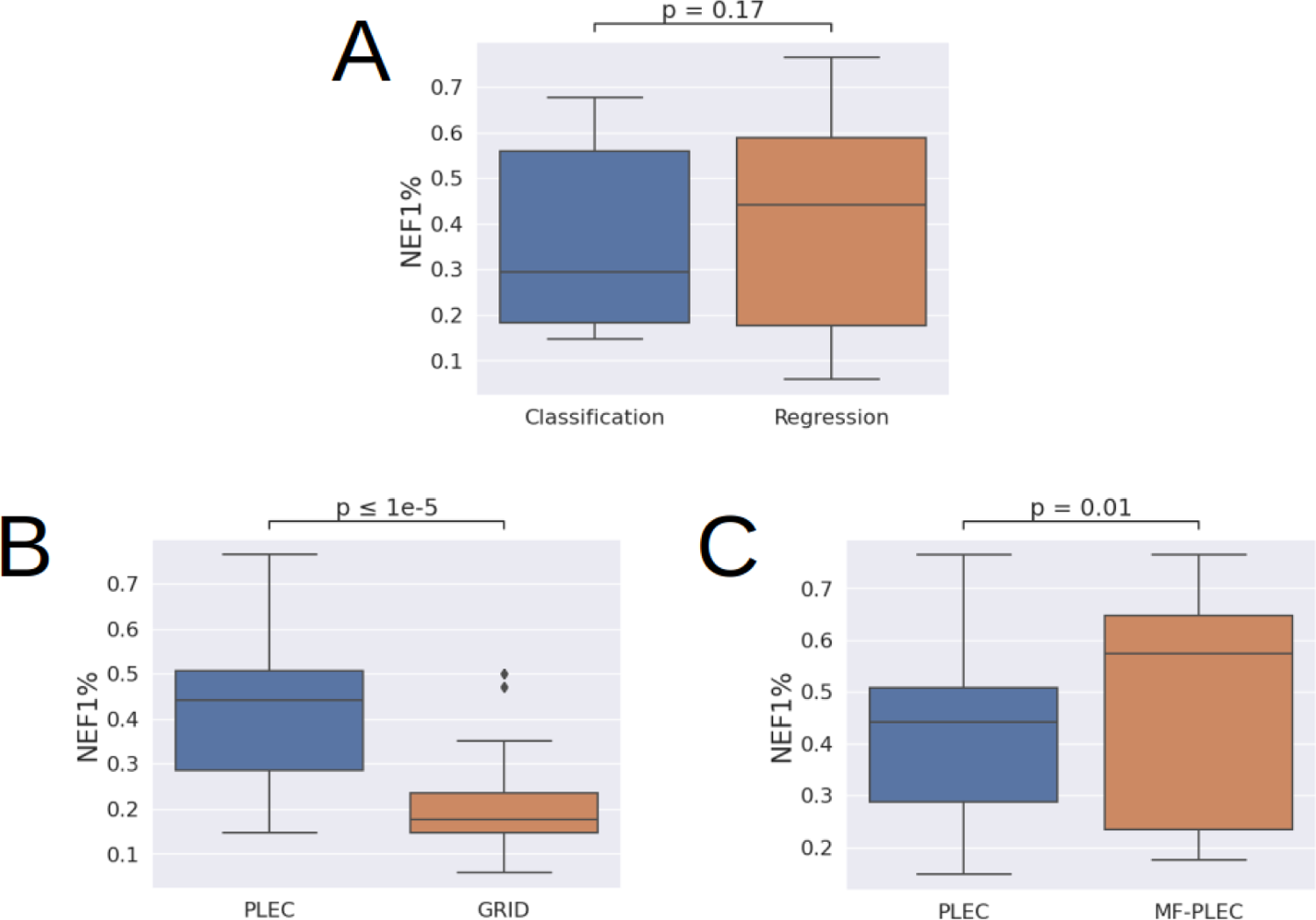
The boxplots illustrating the distributions of NEF1% values according to several computational modeling choices. These include: the nature of the SFs: classification or regression (A); and the featurization scheme: GRID or PLEC (B), PLEC alone or in combination with Morgan fingerprints (C). For each box plot, the median value is represented by a horizontal line inside the box.

The aforementioned best-performing PARP1-specific ML SF, simply referred to as SVM-R from this point, was able to retrieve 26 true active molecules in the top 1% of the SF-ranked test set (Table S4). To assess the diversity in chemical structures of these test actives, we computed their pair-wise Tanimoto similarity in terms of Morgan fingerprints (2048 bits, radius 2), and clustered them accordingly (two compounds with a similarity score ≥ 0.70 are put in the same cluster, as proposed in a recently-developed protocol) [39]. Each of these test actives was also compared to 1565 molecules of the training set, using the same aforementioned Morgan fingerprints: the similarity score to the closest (most similar) training molecule was recorded for each true hit. Results are depicted in Figure 7.

**Figure 7.**
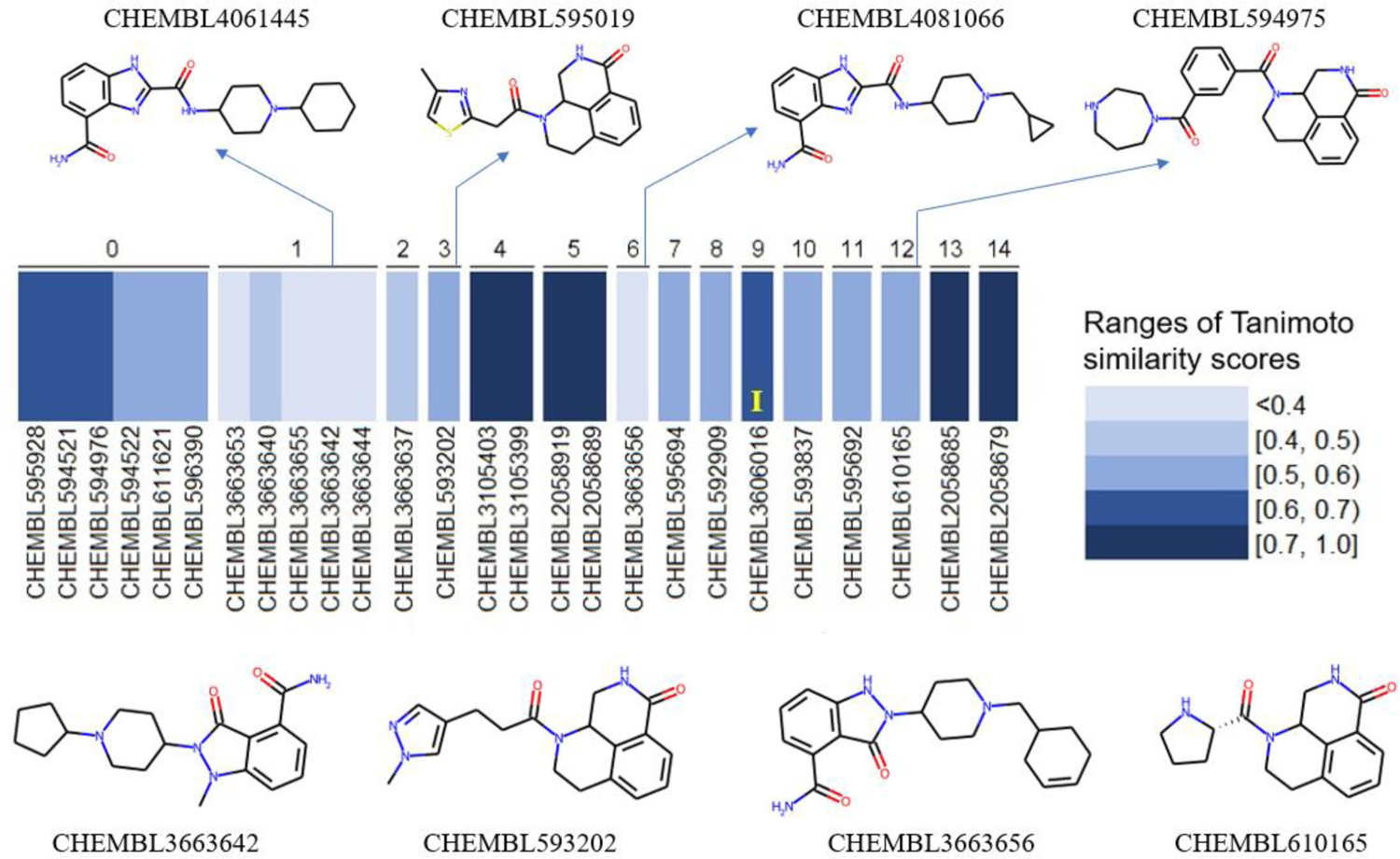
15 clusters according to Tanimoto similarity scores of the 26 test actives retrieved by the SVM-R. For each cluster, a heatmap depicting the Tanimoto similarity score (Morgan fingerprints, 2048 bits, radius 2) between each retrieved true hit and its closest (most similar) training molecule is provided. The structures of several representative exclusive true hits are depicted in 2D (bottom row structures), along with their corresponding closest training molecules (top row structures). The yellow letter “I” in the heatmap area marks the only true hit whose corresponding closest training molecule is an inactive instance.

## Discussion

In this study, state-of-the-art approaches were used to train different ML SFs specific to screening PARP1 inhibitors. Five supervised learning algorithms (each with a binary classification variant and a regression variant) and a Bayesian optimization method for hyperparameter tuning were employed. In a non-systematic preliminary stage (data not shown), we analyzed four PDB structures of this target (PDB IDs: 7AAC, 7KK5, 7KK4 and 6VKK). A systematic analysis was then conducted, focusing on the modeling choices that gave the best performance: 7KK4 as the PARP1 template structure, CNN-Score and SCORCH as the generic ML SFs, and PLEC and GRID as the featurization schemes. This structure-based approach was motivated by comprehensive comparative studies across targets showing that models exploiting protein-ligand features outperform those based on ligand-only features [68,70,71].

The performance of the generic SFs (Smina, CNN-Score, SCORCH) on this target was moderate (Smina) to good (CNN-Score and SCORCH), the two ML-based ones outperformed the classical Smina alone. The improvement of Smina by ML rescoring has also been seen for many other targets [39]. Target-specific ML SFs are generally better than their generic counterparts [39,65]. Therefore, given that Singh *et al* had successfully used a generic classification model trained on inactive-enriched data (chemical features and pharmacophores) to identify PARP1 inhibitors with sub-micromolar activity [72], this makes target-specific ML SFs even more promising for this target [73].

These results also provide yet another example of deep learning not leading to the most predictive model. This has also been observed with other tabular data sets [74–76], which further supports the importance of comprehensive ML analysis. Employing the right featurization strategy for the considered target can also be critical: PLEC-based ML SFs were significantly better-performing than GRID-based ones. While the use of target-specific SFs normally leads to a smaller domain of applicability, the lack of confounding factors related to features-activity relationships may enable more accurate, and therefore better, screening performance. PARP1-specific ML SFs indeed outperformed generic ones in some cases, which is consistent with published reports that suggested the superiority of target-specific models over multi-target ones [77]. Thus, it is worthwhile to spend the resources required to build SFs tailored to PARP1, whose procedure partly involves carefully curating the bioactivity data from several assays, as it was the case here. Often, these ML SFs were far more predictive than non-ML SFs [23,29]. Note that PLEC features, as implemented, do not have direct correspondences to protein-ligand interactions and, hence, the resulting SFs are not amenable to interpreting their predictions, unlike less predictive sets of features [78].

Since there was a strong dependency of SFs used for SBVS on the fundamental physical properties of protein-ligand complexes, we investigated methods to augment the training set to see whether this practice would have an impact on model training and VS performance. We employed data derived from the topological properties of the ligands themselves, in addition to protein-ligand data. We found that the addition of Morgan fingerprint features in the training process enhanced the VS performance for the most PARP1-specific SFs (7 out of 10 models) and was statistically significant (Figure 6C). However, using combined PLEC and Morgan fingerprint features led to no further increase of NEF1% for the best-performing SVM regressor. Two PARP1-specific ML models strongly improved the VS performance offered by generic SFs, as they reached the EF1% of 0.764 on the full test set. These models still outperformed all other SFs when the test molecules were structurally dissimilar to the training molecules. Both models were based on SVM regression, with one utilizing exclusively complex-based PLEC fingerprints. This rendered it notably more efficient than the other model, which incorporated a combination of PLEC and Morgan fingerprint features.

As observed in Figure 7, the true hits retrieved by SVM-R are quite different in terms of chemical structure: out of 15 clusters according to Morgan fingerprints, 13 contain no more than two molecules each (eleven of them each comprise a single active). A comparison of these test actives with all training molecules reveals that most of them are not similar to any molecule of the training set either: most Tanimoto similarity scores (Morgan fingerprints, 2048 bits, radius 2), even to the closest (most similar) training instance, do not exceed 0.60 (16 out of 26, i.e., 61.54%; this percentage is 76.92% if the similarity threshold is 0.70; Table S5). Among these 26 true hits, there is one whose closest training molecule is an inactive instance. This suggests that the performance of our best-performing ML SF was not spoilt by negative nearest neighbors: it could recognize a true active from the test set even though the most structurally similar molecule to this active in the training data is an inactive. Overall, the chemical structures of the test actives retrieved by SVM-R for this target are diverse and cover a large chemical space (even outside that of the training data), which is of particular interest in large-scale prospective VS scenarios where practically all screened molecules will be dissimilar to those in a training set.

## Conclusion

SFs for SBVS are useful to discover novel starting points for the drug discovery process. The development of a high-performing ML SF specific to the investigated target is therefore important and will continue to benefit from artificial intelligence innovation [79]. Here we have seen how much the predictive performance of these SFs against PARP1 varies depending on the featurization scheme, the size and the diversity of available data sets as well as the methodology that is employed. It must be noted that narrow analysis will lead to SFs with suboptimal performance. The SVM-based regressor employing protein-ligand (PLEC) features outperformed all other SFs on both versions (full and dissimilar) of the test set. In particular, rescoring Smina poses with PARP1-specific SFs boosts the retrieval of novel PARP1 inhibitors with respect to using Smina alone. In conclusion, owing to the sufficient availability of experimental and synthetic data instances, a powerful target-specific ML SF has been built and released to carry out SBVS for PARP1 inhibitors.

## Materials and Methods

### Data

PARP1 inhibitors (*n* = 5097) were obtained from the ChEMBL database, version 29 [80]. Only molecules whose bioactivity values were supplied with a standard relation type as “=” (certain data) or “>” (classified as inactives) were kept. To build regression models, each inactive was assigned a pIC_50_ of 2 (we made no claim about the optimality of this choice). The ChemAxon’s Standardizer was used to standardize compounds with the same parameter settings as in a previous study [81]. The average IC_50_ was calculated for a compound if multiple IC_50_ values were available. Redundant compounds (i.e., same ChEMBL ID) with different bioactivity values were kept if the standard deviation of IC_50_s was lower than 2. Compounds with missing SMILES were removed. The bioactivities were annotated based on the bioassay types, including cellular IC_50_s (cell-based assays as a means of primary screening), biochemical IC_50_s (*in vitro* assays against the recombinant PARP1 catalytic domain), and binding affinity values (biophysical assays with the purified PARP1 protein).

The inhibitory activity concentrations of compounds were subject to negative logarithmic transformation: pIC_50_ = -log(IC_50_ x 10^-9^), all IC_50_ values in nanomolar. The training set consists of 1565 molecules with biochemical (but not cellular) IC_50_s: 957 of them have IC_50_s ≤ 1 µM (pIC_50_s ≥ 6), and the other 608 have IC_50_s > 1 µM (pIC_50_s < 6). The test set is composed of 93 molecules annotated with cellular IC_50_s (in addition to *in vitro* biochemical potency): 67 of them are actives (cellular pIC_50_s ≥ 6). Note that, these 67 test actives also have biochemical potency in the active range (≥ 6); while the 26 true test inactives, on the other hand, were determined solely based on their cellular potency (< 6), regardless of their biochemical pIC_50_s, which are typically better than the corresponding cellular values. The test actives were subsequently used as input for DeepCoy to generate property-matched decoys with a decoy-to-active ratio of 50 (giving 3350 decoys in total). These decoys form part of the test set, making it comprise 3443 molecules in total. The pIC_50_s of the molecules in the training and test sets fall into the same range.

A dissimilar test set was generated from the above full test set, composed uniquely of test molecules whose Tanimoto similarity (in terms of Morgan fingerprints, 2048 bits, radius 2) to any training instances was lower than 0.70. This smaller test population thus consists of 3426 molecules (57 true actives, 19 true inactives, 3350 DeepCoy decoys). The code for computing Tanimoto similarity from Morgan fingerprints and all training-test data are provided in our github repository, indicated in the “Data and materials availability” section.

### Selection of protein structures

Although the full-length PARP1 structure has yet to be crystallized or structurally defined [82], the structural characterization of the protein’s interactions with its ligand using its isolated catalytic region appears to be adequate [82]. The catalytic domain of PARP1 includes an alpha-helical N-terminal domain and a mixed alpha/beta C-terminal ADP-ribosyltransferase domain. The co-crystallized ligand of the catalytic domain was shown to bind with the nicotinamide-binding pocket via extensive hydrogen bonds and π-π stacking as well as hydrophobic interactions [82]. Such understanding of the catalytic and inhibitory mechanisms of PARP1 provides an insight into the development of therapeutic agents targeting PARP1.

A total of 78 PARP1 experimental structures were available in the Protein Data Bank (PDB) [83], as of September 2021. Among 19 crystal structures which were of good resolutions (1.50-2.10 Å), we manually selected four PDB IDs: 7AAC, 7KK5, 7KK4, and 6VKK. Each of them was co-crystalized with a small molecule non-covalently bound to PARP1’s catalytic domain. Of these four structures, Val762 was replaced with an alanine in 6VKK and 7AAC. Finally, the catalytic domain of 7KK4 (chain A) [82] was selected to represent the receptor of PARP1 for this study.

### Molecular docking

The Open Babel software [84] was used to generate 3D conformations for all screened compounds using the MMFF94 force field (*--gen3d* option), giving input for the Smina docking software [53]. On the other hand, the DockPrep tool from Chimera [85] was used to prepare the receptor structure (35 amino acids and 4 water molecules) for docking. Partial charges of histidines in the receptor were assigned by the AM1-BBC method [86]. For docking with Smina [53], the search space was centered on the position of the co-crystallized ligand (olaparib of 7KK4), and the size of each axis was set at 30 Å, giving all ligands sufficient space to rotate. Only one pose having the best Smina docking score was retained for each molecule. It is noteworthy that Smina employs a stochastic conformational sampling approach to generate docking poses [87]. Therefore, in principle, there could be significant differences in the best docking pose per molecule if the docking run is repeated. In practice, as each docking run repeats the optimization of a molecule eight times (Smina default value for exhaustiveness setting), the best docking pose per molecule is, on average, stable across runs, and hence, a minimal impact on the results would be observed even when the experiments are repeated. This can be easily checked using the released code.

### Featurization

Each protein-ligand complex was encoded using one of the following featurization strategies. First, 3D grid-based (GRID) features were extracted using the *RDKitGridFeaturizer* function from the *deepchem* Python package [88] with the following options: *ecfp_power* = 9, *splif_power* = 9, *voxel_width* = 16.0, giving 2052 features in total. Second, protein-ligand extended connectivity (PLEC) fingerprints were extracted to describe the interactions between the protein and the ligand using the *PLEC* function from the *ODDT* (Open Drug Discovery Toolkit) Python package [89]. The fragment depth was set at 1 and 5 for the ligand (*depth_ligand*) and the protein (*depth_protein*), respectively, and the fingerprint size was set at 4092, as observed in a past study [51]. Third, Morgan fingerprint features are independent of the protein-ligand complex and its intermolecular interactions (this is the essential difference from GRID and PLEC features) [52]. They were generated using *GetMorganFingerprintAsBitVect* function from the *RDKit* Python package with the following options: *radius = 2, n_bits = 512*. As only the atoms of the ligand are taken into account (no receptor atom is involved), these descriptors are thus receptor-independent, and were chosen to complement protein-ligand descriptors for the featurization of the data sets prior to ML modeling.

### Generic SFs

Three generic SFs were applied to score the docked poses of the test set, including Smina [53], CNN-Score [55] and SCORCH [54]. Relevant information on these three SFs can be found in the “Applying existing generic SFs” part of the “Results” section.

### Construction of PARP1-specific ML SFs

Several ML algorithms including random forest (RF) [46], extreme gradient boosting (XGB) [47], support vector machine (SVM) [48], artificial neural network (ANN) [49], and deep neural network (DNN) [50] were used for both classification and regression models.

RF and XGB are both ensemble models composed of decision trees (DTs). However, their working principles are different, in that the former is based on bootstrap aggregation, whereas the latter functions on the boosting principle. Indeed, each DT in an RF is trained independently, after which their predictions on new data are decided either by majority voting (in case of classifiers), or by averaging individual output (in case of regressors). XGB, on the other hand, comprises sequential DTs, i.e., they are trained in succession, such that the errors committed by earlier DTs are corrected or minimized by later ones. Another traditional ML algorithm used in this study is SVM, which seeks to construct a hyperplane that separates the data points representing all data instances, in a way that this plane is the furthest possible to its nearest data point. Besides, two algorithms inspired by the human brain, ANN and DNN, are employed to train our models. Their main components are an input layer, one or more hidden layers (a DNN is an ANN with at least two hidden layers), and an output layer. Multiple neurons constitute the hidden nodes, and are responsible for all the computations that take place after the input data are provided.

All ML procedures were carried out using the *sklearn* [90], *XGBoost* [47], and *keras* python packages [90]. They were used to develop target-specific ML SFs and hyperparameters were optimized with the *hyperopt* package. Our Python code and details regarding the hyperparameters are provided at https://github.com/cabaklaud/SBVS-PARP1. The VS performance of the models (both classifiers and regressors) with optimal hyperparameters was evaluated.

### Hyperparameter tuning

ML algorithms are increasingly used in SBVS, owing to their robust performance. Hyperparameter optimization may lead to improvements in the predictive performance as different data have their own unique characteristics. A poorly-configured algorithm may not perform better than a random-guessing model, while a well-configured one may achieve good prediction accuracy. Bayesian optimization is an efficient method to optimize the hyperparameters of an ML algorithm. It is based on a sequential search framework that incorporates both exploration and exploitation. In this study, we used the *hyperopt* Python library to search for optimal hyperparameters for model building [91]. There are three factors that must be defined: the search space, the objective function, and the optimization algorithm. The search space is the space of hyperparameters and their values, which can be defined by users. The objective function computes the statistical assessment errors from the five-fold cross-validation of ML algorithms using all training data. The optimization algorithm is a model that compares various hyperparameter values within the search space and finds the optimal one to minimize the loss, for example, the Tree-structured Parzen Estimator (TPE). The TPE is a method that builds models to predict the performance of hyperparameters based on historical measurements, and then subsequently chooses new hyperparameters. The *fmin* function from *hyperopt* was used to carry out the optimization process by minimizing the function over a given configuration space, storing the results and finding the best-performing configuration of hyperparameters. Bayesian optimization was used to search for the best values of the hyperparameters of each algorithm within the provided range (Table S5).

### Measuring virtual screening performance

In this study, we calculated the enrichment factor at the top 1%-ranked molecules (EF1%) [56], and the normalized EF1% (NEF1%) [57] to evaluate VS performance. The EF1% and NEF1% values were computed using an in-house bash script published as part of a recent protocol [39]. The *metrics.precision_recall_curve* function of the *sklearn* Python library was used to compute PR values and plot PR curves [90].

## Abbreviations

ANN: Artificial neural network

ART: ADP-ribosyl transferase

BRCA1: Breast Cancer Type 1 Susceptibility Protein

BRCA2: Breast Cancer Type 2 Susceptibility Protein

BRCT: BRCA1 C-terminus domain

CAT: Catalytic domain

CNN: Convolutional neural network

DNN: Deep neural network

DUD-E: Database of Useful Decoys: Enhanced

EF1%: Enrichment Factor at the top 1%

HD: Helical domain

ML: Machine learning

MMEJ: Microhomology-mediated end joining

NEF1%: Normalized Enrichment Factor at the top 1%

PARP1: Poly ADP-ribose polymerase 1

RF: Random forest

SB: Structure-based

SF: Scoring function

SVM: Support vector machine

VS: Virtual screening

XGB: Extreme gradient boosting

TPE: Tree-structured Parzen Estimator

## Acknowledgments

Funding: This work was supported by the Foundation ARC pour la Recherche sur le Cancer [ARC to V.K.T.N.]; the Wolfson Foundation and the Royal Society for a Royal Society Wolfson Fellowship [to P.J.B.].

## Author contributions

Conceptualization: T.R., P.J.B.

Methodology: K.C., V.K.T.N., P.J.B.

Investigation: K.C., V.K.T.N., P.J.B.

Visualization: K.C., V.K.T.N., P.J.B.; Supervision: P.J.B.

Writing—original draft: V.K.T.N., P.J.B.

Writing—review & editing: K.C., V.K.T.N., T.R., P.J.B.

## Other acknowledgments

We are also thankful to the developers of the employed software packages (ChemAxon, OpenBabel, Gnina, Chimera, Scikit-learn). Thanks also to Muhammad Junaid and Saw Simeon for their feedback on this work.

## Data and materials availability

Code and data sets which can be used to reproduce the results in this paper are freely available at https://github.com/cabaklaud/SBVS-PARP1.

