## supplementary materials for "Comprehensive machine learning boosts structure-based virtual screening for PARP1 inhibitors"

**Table S1.** Performance on the full test set of three generic SFs and PARP1-specific ML SFs using protein-ligand features (PLEC and GRID). For each target-specific ML SF, the EF1% and NEF1% indicated herein are median values obtained after 10 training-test runs. For each generic SF, only one test run was performed. The maximal EF1% is 50.75.

| **Scoring function** | **Method** | **Type** | **EF1%** | **NEF1%** |
|---|---|---|---|---|
| Smina | Generic | Regression | 11.94 | 0.235 |
| CNN-Score | Generic | Classification/Regression | 19.40 | 0.382 |
| SCORCH | Generic | Classification | 17.91 | 0.352 |
| PLEC RF | PARP1-specific | Classification | 26.115 | 0.5145 |
| PLEC XGB | PARP1-specific | Classification | 20.89 | 0.411 |
| PLEC SVM | PARP1-specific | Classification | 32.83 | 0.647 |
| PLEC ANN | PARP1-specific | Classification | 10.44 | 0.205 |
| PLEC DNN | PARP1-specific | Classification | 13.43 | 0.264 |
| GRID RF | PARP1-specific | Classification | 8.95 | 0.176 |
| GRID XGB | PARP1-specific | Classification | 8.95 | 0.176 |
| GRID SVM | PARP1-specific | Classification | 25.37 | 0.5 |
| GRID ANN | PARP1-specific | Classification | 8.95 | 0.176 |
| GRID DNN | PARP1-specific | Classification | 11.94 | 0.235 |
| PLEC RF | PARP1-specific | Regression | 23.88 | 0.470 |
| PLEC XGB | PARP1-specific | Regression | 22.38 | 0.441 |
| PLEC SVM | PARP1-specific | Regression | 38.8 | 0.764 |
| PLEC ANN | PARP1-specific | Regression | 15.665 | 0.3085 |
| PLEC DNN | PARP1-specific | Regression | 13.43 | 0.2645 |
| GRID RF | PARP1-specific | Regression | 7.46 | 0.147 |
| GRID XGB | PARP1-specific | Regression | 7.46 | 0.147 |
| GRID SVM | PARP1-specific | Regression | 23.88 | 0.47 |
| GRID ANN | PARP1-specific | Regression | 7.46 | 0.147 |
| GRID DNN | PARP1-specific | Regression | 8.95 | 0.176 |
| Expected if classification is carried out at random | | | 1.00 | 0.020 |

**Table S2.** Performance on the full test set of PARP1-specific ML SFs using ligand-only features (Morgan fingerprints) combined with protein-ligand features (PLEC). For each target-specific ML SF, the EF1% and NEF1% indicated herein are median values obtained after 10 training-test runs. The maximal EF1% is 50.75.

| **Scoring function** | **Method** | **Type** | **EF1%** | **NEF1%** |
|---|---|---|---|---|
| Morgan fingerprint + PLEC RF | PARP1-specific | Classification | 32.83 | 0.647 |
| Morgan fingerprint + PLEC XGB | PARP1-specific | Classification | 28.35 | 0.558 |
| Morgan fingerprint + PLEC SVM | PARP1-specific | Classification | 34.32 | 0.676 |
| Morgan fingerprint + PLEC ANN | PARP1-specific | Classification | 10.44 | 0.205 |
| Morgan fingerprint + PLEC DNN | PARP1-specific | Classification | 10.44 | 0.205 |
| Morgan fingerprint + PLEC RF | PARP1-specific | Regression | 32.83 | 0.647 |
| Morgan fingerprint + PLEC XGB | PARP1-specific | Regression | 29.85 | 0.588 |
| Morgan fingerprint + PLEC SVM | PARP1-specific | Regression | 38.8 | 0.764 |
| Morgan fingerprint + PLEC ANN | PARP1-specific | Regression | 18.65 | 0.367 |
| Morgan fingerprint + PLEC DNN | PARP1-specific | Regression | 20.89 | 0.4115 |
| Expected if classification is carried out at random | | | 1.00 | 0.020 |

**Table S3.** Performance on the dissimilar test set of three generic SFs and PARP1-specific ML SFs using protein-ligand features (PLEC) or a combination of ligand-only and protein-ligand features (Morgan fingerprint + PLEC). The dissimilar test set includes 57 true actives and 3369 inactives (19 true inactives and 3350 DeepCoy-generated decoys). For each target-specific ML SF, the EF1% and NEF1% indicated herein are median values obtained after 10 training-test runs. For each generic SF, only one test run was performed. The maximal EF1% is 59.65.

| **Scoring function** | **Method** | **Type** | **EF1%** | **NEF1%** |
|---|---|---|---|---|
| Smina | Generic | Regression | 7.01 | 0.117 |
| CNN-Score | Generic | Classification/Regression | 17.54 | 0.294 |
| SCORCH | Generic | Classification | 22.80 | 0.382 |
| PLEC RF | PARP1-specific | Classification | 21.05 | 0.352 |
| PLEC XGB | PARP1-specific | Classification | 14.03 | 0.235 |
| PLEC SVM | PARP1-specific | Classification | 35.08 | 0.588 |
| PLEC ANN | PARP1-specific | Classification | 0 | 0 |
| PLEC DNN | PARP1-specific | Classification | 7.01 | 0.117 |
| Morgan fingerprint + PLEC RF | PARP1-specific | Classification | 31.57 | 0.529 |
| Morgan fingerprint + PLEC XGB | PARP1-specific | Classification | 21.05 | 0.352 |
| Morgan fingerprint + PLEC SVM | PARP1-specific | Classification | 35.08 | 0.588 |
| Morgan fingerprint + PLEC ANN | PARP1-specific | Classification | 0 | 0 |
| Morgan fingerprint + PLEC DNN | PARP1-specific | Classification | 0 | 0 |
| PLEC RF | PARP1-specific | Regression | 22.80 | 0.382 |
| PLEC XGB | PARP1-specific | Regression | 19.29 | 0.323 |
| PLEC SVM | PARP1-specific | Regression | 35.08 | 0.588 |
| PLEC ANN | PARP1-specific | Regression | 8.77 | 0.147 |
| PLEC DNN | PARP1-specific | Regression | 7.89 | 0.132 |
| Morgan fingerprint + PLEC RF | PARP1-specific | Regression | 34.205 | 0.573 |
| Morgan fingerprint + PLEC XGB | PARP1-specific | Regression | 24.56 | 0.411 |
| Morgan fingerprint + PLEC SVM | PARP1-specific | Regression | 35.08 | 0.588 |
| Morgan fingerprint + PLEC ANN | PARP1-specific | Regression | 12.275 | 0.2055 |
| Morgan fingerprint + PLEC DNN | PARP1-specific | Regression | 12.275 | 0.2055 |
| Expected if classification is carried out at random | | | 1.00 | 0.017 |

**Table S4.** True active molecules retrieved in the top 1% of the full test set by the best-performing PARP1-specific ML SFs SVM-R. For each retrieved true active, the most similar (closest) training molecule and training active (in terms of Morgan fingerprints, 2048 bits, radius 2) are identified (same molecule in most cases), and the corresponding Tanimoto similarity score is computed. The potency ($pIC_{50}$) reported in ChEMBL of each true hit is also provided. The closest training molecule for all test actives was an active except for CHEMBL3606016, for which it was an inactive. Its closest training active was CHEMBL4081066 ($pIC_{50}$ = 6.97, Tanimoto similarity = 0.285714286).

| Test actives | Potency ($pIC_{50}$) of the test active | Closest training molecule | Potency ($pIC_{50}$) of the closest training molecule | Similarity to the closest training molecule |
|---|---|---|---|---|
| CHEMBL595928 | 6.8 | CHEMBL595019 | 8.22 | 0.615384615 |
| CHEMBL594521 | 6.7 | CHEMBL595019 | 8.22 | 0.619047619 |
| CHEMBL594976 | 6.19 | CHEMBL595019 | 8.22 | 0.619047619 |
| CHEMBL594522 | 6.24 | CHEMBL595019 | 8.22 | 0.557142857 |
| CHEMBL611621 | 6.7 | CHEMBL595019 | 8.22 | 0.590909091 |
| CHEMBL596390 | 6.46 | CHEMBL595019 | 8.22 | 0.578125 |
| CHEMBL3663653 | 6.72 | CHEMBL4081066 | 6.97 | 0.375 |
| CHEMBL3663640 | 7.58 | CHEMBL4061445 | 6.51 | 0.411764706 |
| CHEMBL3663655 | 7.36 | CHEMBL4081066 | 6.97 | 0.342105263 |
| CHEMBL3663642 | 6.89 | CHEMBL4061445 | 6.51 | 0.397058824 |
| CHEMBL3663644 | 7.3 | CHEMBL131737 | 7.5 | 0.355932203 |
| CHEMBL3663637 | 6.62 | CHEMBL4061445 | 6.51 | 0.461538462 |
| CHEMBL593202 | 6.38 | CHEMBL595019 | 8.22 | 0.565217391 |
| CHEMBL3105403 | 6.06 | CHEMBL3105405 | 7.25 | 0.701754386 |
| CHEMBL3105399 | 6.15 | CHEMBL3105400 | 6.96 | 0.918367347 |
| CHEMBL2058919 | 8.22 | CHEMBL3903698 | 8.89 | 0.859375 |
| CHEMBL2058689 | 6.91 | CHEMBL3981021 | 6.6 | 0.830188679 |
| CHEMBL3663656 | 6.82 | CHEMBL4081066 | 6.97 | 0.391891892 |
| CHEMBL595694 | 6.89 | CHEMBL595019 | 8.22 | 0.52238806 |
| CHEMBL592909 | 6.51 | CHEMBL594975 | 7.3 | 0.550724638 |
| CHEMBL3606016 | 7 | CHEMBL3605999 | 5 | 0.602941176 |
| CHEMBL593837 | 6.26 | CHEMBL594975 | 7.3 | 0.555555556 |
| CHEMBL595692 | 6.16 | CHEMBL595019 | 8.22 | 0.53030303 |
| CHEMBL610165 | 6.75 | CHEMBL594975 | 7.3 | 0.569230769 |
| CHEMBL2058685 | 6.87 | CHEMBL3903759 | 8.3 | 0.806451613 |
| CHEMBL2058679 | 7.96 | CHEMBL2058680 | 8.52 | 0.907407407 |

**Table S5.** A detailed list of search spaces used for Bayesian optimization. Letters C and R preceding the search space indicate whether the search space was used for classification or regression models. The absence of the letter means the same search space was used for both regression and classification-based models.

| Algorithm | Parameter | Expression | Search space |
|---|---|---|---|
| DNN (8 layers) | Neurons per layer | Uniform | {64:8192} |
| | Batch size | Uniform | {128:500} |
| | Activation function | Choice | {‘relu’} |
| | Optimization method | Choice | {Adam,Adadelta,RMSprop} |
| | Dropout rate | Uniform | {0:1} |
| RF | Number of trees in the forest | Uniform | {100:10000} |
| | Maximum depth of the trees | Choice | {1, 2, 3, 4, 5, None} |
| | Function to measure the quality of a split | Choice | C: {'gini', 'entropy’}<br>R: {'squared_error', 'absolute_error', 'friedman_mse', 'poisson'} |
| SVM | Regularization parameter | Uniform | {0:20} |
| | Kernel coefficient | Choice | {‘scale’, ‘auto’} |
| | Kernel type | Choice | {‘rbf’, ‘poly’, ‘sigmoid’} |
| ANN | The number of neurons in the hidden layer | Uniform | {8:140} |
| | Activation function for the hidden layer | Choice | {‘relu’,’tanh’} |
| | Maximum number of iterations | Uniform | {1000:10000} |
| XGB | Maximum tree depth for base learner | Uniform | {3:18} |
| | Minimum loss reduction | Uniform | {1:9} |
| | L1 regularization term on weights | Uniform | {40:180} |
| | L2 regularization term on weights | Uniform | {0:1} |
| | Subsample ratio of columns when constructing each tree | Uniform | {0.5:1} |
| | Minimum sum of instance weights | Uniform | {0:10} |
| | Number of gradient boosted trees | Uniform | {1000:5000} |
